# Network synchronization and synchrony propagation: emergent elements of inspiration

**DOI:** 10.1101/664946

**Authors:** Sufyan Ashhad, Jack L Feldman

## Abstract

The preBötzinger Complex (preBötC) – the kernel of breathing rhythmogenesis in mammals – is a non-canonical central pattern generator with undetermined mechanisms. We assessed preBötC network dynamics under respiratory rhythmic and nonrhythmic conditions *in vitro*. In each cycle under rhythmic conditions, an inspiratory burst emerges as (presumptive) preBötC rhythmogenic neurons transition from aperiodic uncorrelated population spike activity to become increasingly synchronized during preinspiration, triggering bursts; burst activity subsides and the cycle repeats. In a brainstem slice in nonrhythmic conditions, antagonizing GABA_A_ receptors can initiate this periodic synchronization and consequent rhythm coincident with inducing a higher conductance state in nonrhythmogenic preBötC output neurons. Furthermore, when input synchrony onto these neurons was weak, preBötC activity failed to propagate to motor nerves. Our analyses uncover a dynamic reorganization of preBötC network activity – underpinning intricate cyclic neuronal interactions leading to network synchronization and its efficient propagation – correlated with and, we postulate, essential to, rhythmicity.

## INTRODUCTION

While now well established that the preBötC is the rhythmogenic kernel for breathing (Smith et al., 1991), the underlying mechanism(s) have eluded discovery (Del Negro et al., 2018). Critically, two long-favored hypotheses for rhythmogenesis, i.e., that rhythm is generated by pacemaker neurons or by simple circuits dependent on inhibition including postinhibitory rebound, are not supported by experimental tests (Baertsch et al., 2018; Del Negro et al., 2018; Feldman et al., 2013; Feldman and Kam, 2015; Janczewski et al., 2013; Rekling and Feldman, 1998; Sherman et al., 2015). Here, we explore an alternative (though nonexclusive) hypothesis for preBötC rhythmogenesis, that the rhythm is an emergent property of the preBötC microcircuit (Del Negro et al., 2018; Feldman and Kam, 2015; Kam et al., 2013a; Kam et al., 2013b).

Acute slices from neonatal brainstem of rodents containing the preBötC bathed in artificial cerebrospinal fluid (ACSF) with near physiological brain [K^+^] (3 mM) and [Ca^2+^] (1.5 mM) are nonrhythmic (Smith et al., 1991) (Figures 1A, S1A). However, with elevated [K^+^]_ACSF_ (9 mM), rhythmic inspiratory bursts (I-bursts) emerge in preBötC that in almost all (≳90%) instances cause motor-related I-bursts in hypoglossal nerve (XIIn; Figures 1A2-A3, S1A2-A3). In each cycle, for up to a few hundred milliseconds prior to the rapid onset of I-bursts, preBötC population activity grows slowly; this epoch is designated preinspiration (preI) and has a duration of ~250 msec *in vitro* (Figure 1A4). Notably, when preI fails to induce a preBötC I-burst, the result is a low amplitude preBötC burstlet with no associated XIIn I-burst (Kam et al., 2013a). To understand rhythmogenic mechanisms underlying breathing, we addressed several key questions: What kind of network dynamics lead to and underlie preI activity? What network mechanism(s) determine the threshold for transition of preI activity into I-bursts? Which neuronal subpopulations of the preBötC microcircuit contribute to preI and I-burst activity?

**Figure 1.**
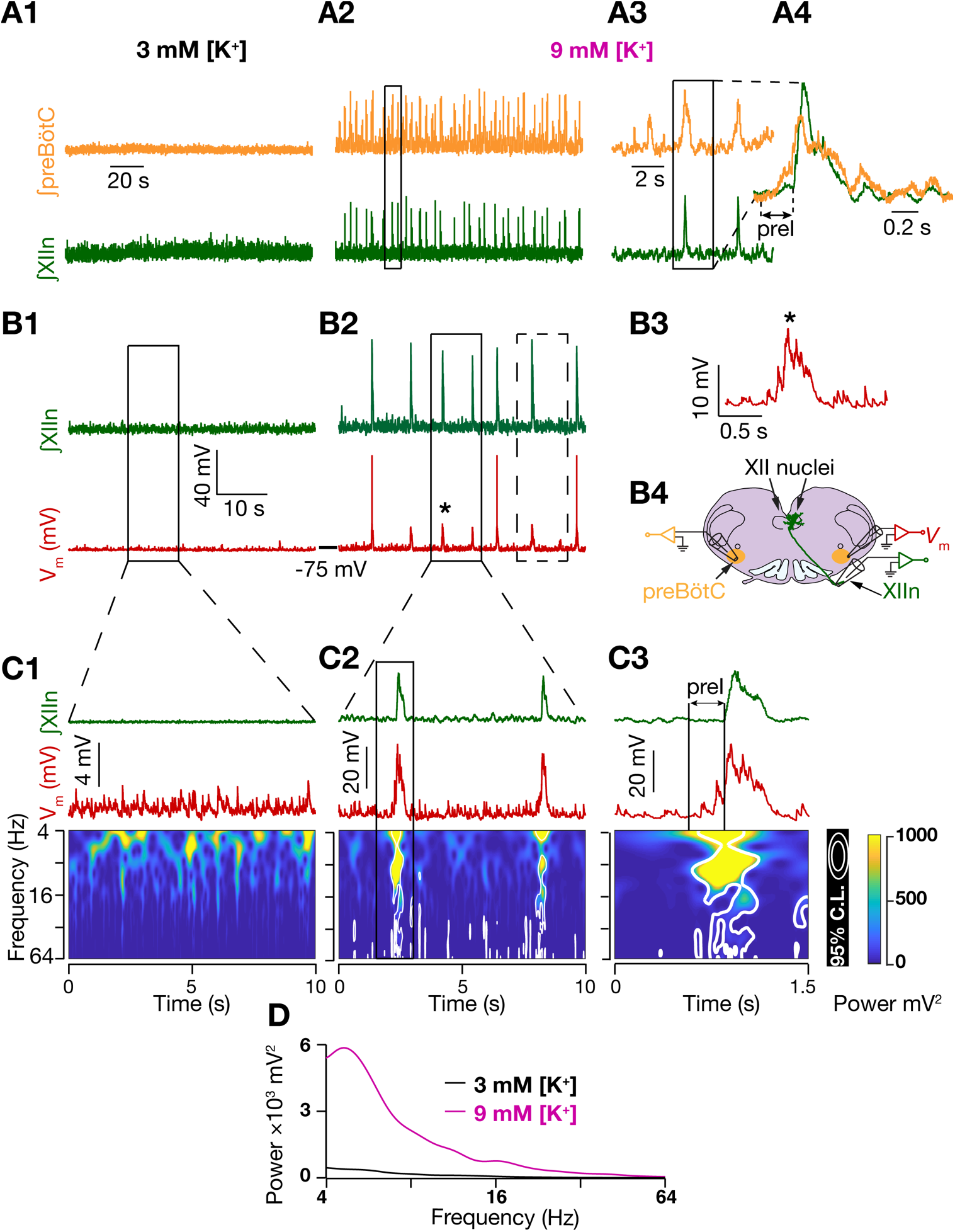
Inputs onto preBötC I-M SST^+^ neurons undergo spectrotemporal reorganization correlated with emergence of inspiratory rhythm. (A) preBötC population activity (orange: integrated preBötC (∫preBötC)); green: integrated XIIn (∫XIIn)) in 3 mM [K^+^]_ACSF_, i.e., control (A1) and in 9 mM [K^+^]_ACSF_ (A2); (A3) expanded boxed region from (A2). (A4) expanded boxed region from (A3) with ∫preBötC and ∫XIIn I-burst overlaid. (B1-B2) ∫XIIn and I-M SST^+^ neuron membrane potential (*V*_m_; red) in 3 mM [K^+^]_ACSF_ (B1) and in 9 mM [K^+^]_ACSF_ (B2); (B3) expanded *V*_m_ from (B2) marked by *; (B4) Configuration for simultaneous recording of SST^+^ neurons in preBötC, XIIn, and contralateral preBötC population activity. preBötC neurons project to XII premotor and motor neurons in the slice. (C1-C3) ∫XIIn, *V*_m_ and associated frequency-time plot for I-M SST^+^ neuron in 3 mM [K^+^]_ACSF_ (nonrhythmic; (C1);*V*_m_ from solid boxed region in (B1)) and in 9 mM [K^+^]_ACSF_ (rhythmic, (C2) *V*_m_ from solid boxed region in (B2)). (C3) expanded boxed region from (C2). White contours in frequency-time plots enclose regions where local power was significantly higher (95% confidence level) than the background spectrum, i.e., global wavelet spectrum of *V*_m_ in (C1)). Simultaneous ∫XIIn activity is also plotted at the top. Note in (C3) emergence of input synchrony in preI period well before the emergence of I-burst in XIIn. (D) global wavelet spectrum of *V*_m_ in (C1) and (C2).

We used genetic labeling to fluorescently tag a molecularly-defined neuronal subpopulation, i.e., somatostatin-expressing (SST^+^) neurons, within the preBötC in mice. These glutamatergic preBötC output neurons project to inspiratory premotor neurons (Cui et al., 2016; Stornetta et al., 2003; Tan et al., 2010; Yang and Feldman, 2018) and are necessary for breathing *in vivo* (Tan et al., 2008). We explored the network dynamics underlying the generation and propagation of preI and I-bursts by recording synaptic inputs impinging upon preBötC SST^+^ neurons as they transitioned from nonrhythmic, i.e., control, to rhythmic conditions. Time-frequency decomposition of *V*_m_ of inspiratory-modulated (I-M) SST^+^ neurons revealed a spectrotemporal reorganization of their synaptic inputs where they transitioned from randomly tonic in the nonrhythmic condition to periodically synchronized in the rhythmic condition. Furthermore, comparison of simultaneous whole-cell current clamp recordings between pairs of I-M SST^+^ neurons with preBötC and downstream hypoglossal population outputs (quadruple electrode recordings) revealed input synchrony onto the recorded pairs was accompanied by short-latency correlated EPSPs during preI and I-burst. This suggests that active network synchronization underlies initiation and maintenance of preI and subsequent emergence of preBötC I-bursts. Exploring network mechanisms underlying synchronization, we uncovered a shift in the excitation-inhibition balance of the network towards excitation that promotes preBötC synchronization. Furthermore, by assessing connectivity profiles preBötC SST^+^ neurons together with pharmacological studies, spectrotemporal analyses and computational modeling, we argue that it is not the overall level of synaptic inputs (presynaptic activity) but a switch from asynchronous to synchronous activity that is hallmark of preBötC rhythmogenesis and I-burst initiation. Our analyses uncover a non-canonical mechanism for the operation of the breathing central pattern generation (bCPG) that reflects intricate interactions among various neuronal subtypes leading to network assembly underlying breathing rhythmicity.

## RESULTS

### Inputs onto preBötC I-M SST^+^ neurons undergo spectrotemporal reorganization correlated with emergence of inspiratory activity

Under nonrhythmic conditions where preBötC population bursts were absent or sporadic (3 mM [K^+^]_ACSF_; Figure 1A1, B1), preBötC SST^+^ neurons received asynchronous synaptic inputs that rarely summated to produce action potentials (APs; Figure 1B1, C1). However, with further depolarization induced by increasing [K^+^]_ACSF_ to 9 mM, their excitatory postsynaptic potentials (EPSPs) became periodic and clustered, producing more APs at shorter intervals; this activity increased slowly at first, ultimately resulting in I-bursts (Figures 1B2-B3; S1B1-B2). Using wavelet analysis to ascertain the EPSP spectrotemporal structure, we found that the membrane potential (*V*_m_) power in the frequency range 4-64 Hz of preBötC I-M SST^+^ neurons (15/29) significantly increased during preI and subsequent I-bursts (Figure 1C1-C2). During this period, synchronization of EPSPs progressively increased, with inputs at lower frequencies appearing first during preI (Figures 1C3, 2F). Two notable points: i) preBötC I-M SST^+^ neurons rarely fired APs during preI (Figure S1B1-B2; similar to inspiratory Type-2 neurons (Rekling et al., 2000)) making them an unlikely source for preI activity, and; ii) the frequency of synaptic inputs in the interburst intervals (IBIs) preceding each preI decreased as compared to that during nonrhythmic conditions where low frequency inputs (4Hz–8 Hz) were only sporadically present (Figure 1C1).

**Figure 2.**
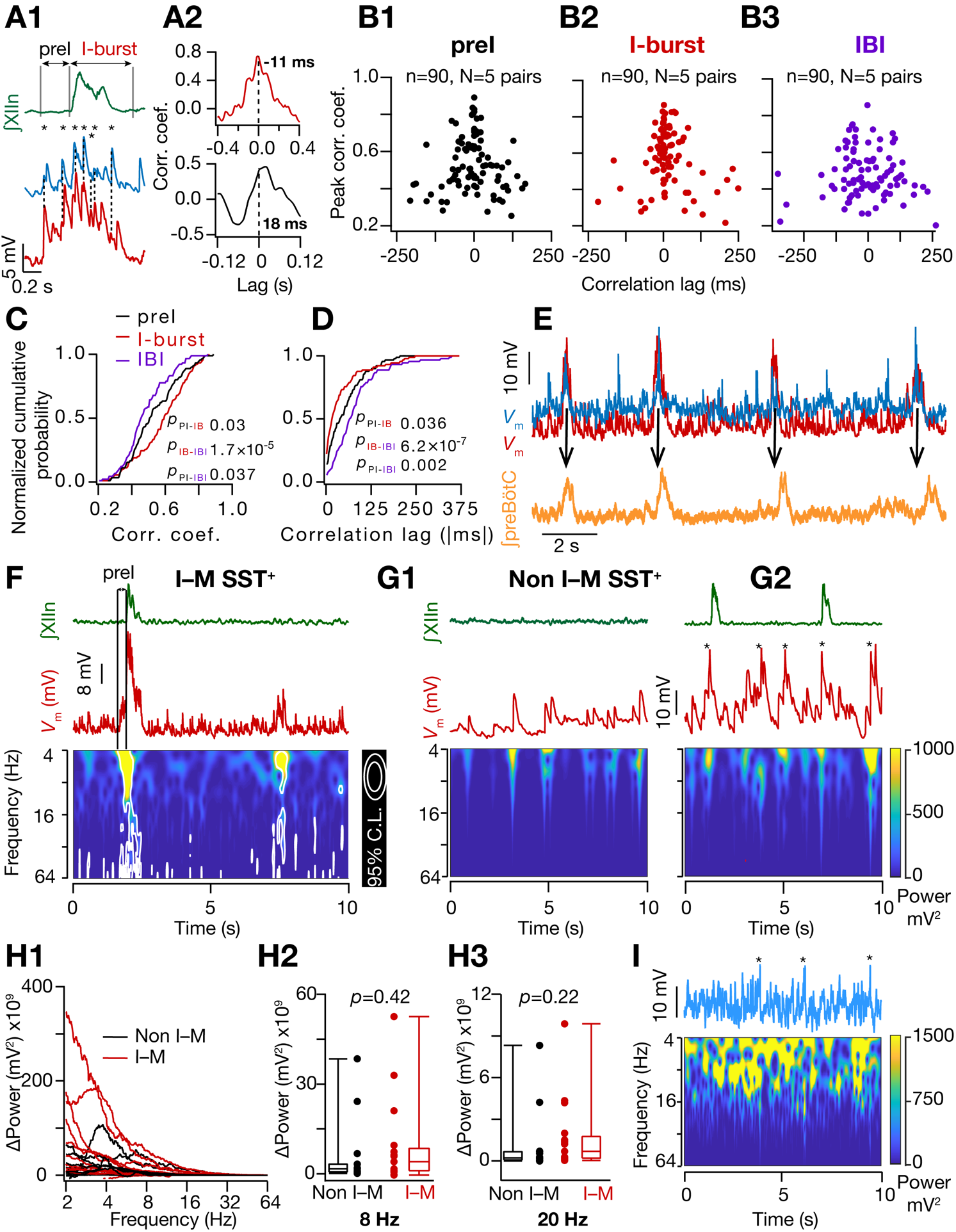
Input synchrony in I-M SST^+^ neurons is associated with increased synaptic correlation between neuron pairs during preI and I-bursts. (A1) ∫XIIn (green) and *V*_m_ of two simultaneously recorded I-M SST^+^ neurons (blue, red) along with the *V*_m_ crosscorrelograms (A2) during preI (black) and I-burst (red). Temporally aligned EPSP peaks indicated by dashed line and *. (B1-B3) Plots for peak correlation vs time lag for *V*_m_ of 5 I-M SST^+^ pairs (90 cycles from 5 pairs) during preI, I-burst and interburst interval (IBI) epochs. (C-D) normalized cumulative histogram of crosscorrelation peaks (C) and lags (D) for events in (B1-B3). Kruskal Wallis test (C, *p*= 5×10^−5^; D, *p*= 9×10^−8^) followed by Wilcoxon signed rank test for pairwise comparisons (*p* values for color-coded pairwise comparisons). (E) *V*_m_ of simultaneously recorded I-M SST^+^ pair (blue, red) along with ∫preBötC (orange) showing that ∫preBötC activity peaks after the peak of *V*_m_ in each cycle. (F) frequency-time plot of *V*_m_ of an I-M SST^+^ neuron (from dashed box region of Figure1(B2)) under rhythmic condition showing reduced synchrony during failure compared to production of I–burst. (G1-G2) same as (F) but for non I-M SST^+^ neuron under control (G1) and rhythmic (9mM [K^+^]_ACSF_) (G2) conditions. *V*_m_ median filtered to remove APs (indicated by *). (H1-H3) change in *V*_m_ power of I- and non-I-M SST^+^ neurons when brainstem slices were shifted from nonrhythmic to rhythmic conditions; *p* values for Wilcoxon rank sum test. (I) frequency-time plot of median filtered *V*_m_ of a model neuron when 10 excitatory synapses were activated randomly with 15 Hz mean frequency. * indicate filtered APs.

### Input synchrony among I-M SST^+^ neurons is concurrent with increased synaptic correlation between neuron pairs during preI and I-bursts

What kind of network interactions result in the emergence of synchronous inputs onto I-M SST^+^ neurons? We hypothesized that these interactions are a consequence of recurrent excitatory connections among these I-M SST^+^ neurons (Guzman et al., 2016; Miles and Wong, 1986). This did not appear to be the case, however, since only 2% (1/50) pairs of preBötC SST^+^ neurons tested were synaptically coupled (Figure S2A). Hence we postulated that the input synchrony of these neurons resulted from convergent inputs arising from afferent rhythmogenic preBötC neurons, consistent with our previous hypothesis (Cui et al., 2016). Putatively rhythmogenic inspiratory preBötC Type-1 neurons (SST^−^) interact through excitatory synapses with ~16% probability of any pair having a one-way connection (Rekling et al., 2000). Notably, in each cycle Type-1 neurons exhibit a progressive slow increase in their firing rate that starts in preI and continues through the I-burst (Gray et al., 1999). In each cycle among pairs of simultaneously recorded I-M SST^+^ neurons, a significant increase in their *V*_m_ correlation was associated with the emergence of preI input synchrony (Figures 2A1-E, S2B1-C3). Importantly, the increase in the *V*_m_ correlation was a consequence of short-latency correlated EPSPs during preI and I-bursts (8/90, 23/90 and 35/90 events had <10 ms correlation lags in the IBI, preI and I-burst epochs respectively; Figures 2B1-B3, D, S2C1-C3); this indicates that input synchrony onto I-M SST^+^ neurons results from synchronization within the preBötC subnetwork generating inspiratory rhythm. Thus, the emergence of periodic synchronous EPSPs in I-M SST^+^ neurons (Figure 1C2) represents a significant spectrotemporal reorganization in the activity of their input population of (presumptive) rhythmogenic neurons. This change in the input structure of I-M SST^+^ neurons only emerges under rhythmic conditions, suggesting that network synchronization is causal to both rhythmogenesis and I-burst generation. Accordingly, when the input synchrony (which we use as a proxy for network synchrony henceforth) onto these neurons was present but weak, there was no resultant I-burst and consequently longer intervals between XIIn bursts (Figures 2F, 4B-C). Furthermore, the increase in the membrane potential power, which signifies an increase in the average frequency of inputs, in non-I-M SST^+^ neurons (14/29 neurons; Figures 2G1-H3, S1C1-C2) was indistinguishable from that of I-M SST^+^ neurons (Figure 2H1-H3), despite the fact that they did not show input synchrony, i.e., significant increases in spectrotemporal power following periods of uncorrelated activity (compare Figure 2F with 2G1-G2; S2D1-D2). This further demonstrates synchronized inputs as a necessary condition for the substantial increase in I-M SST^+^ neuronal activity underlying I-bursts.

We considered the possibility that this synchrony was epiphenomenal, resulting from increased firing rates of individual afferent neurons such that random alignment of more APs appears as synchronous input. We tested this computationally in a biophysically realistic model neuron that had inputs from 10 afferent neurons, each with a Poisson-distributed 2-20 Hz mean AP frequency (See METHODS – *Simulations*). We found no indication of any spectrotemporal reorganization from uncorrelated activity to significant input synchrony (Figure 2I) and no rhythmic bursting (Figures 2I, S2E). We conclude that synchrony is not the result of an increase in network excitability but is a manifestation of network assembly (Buzsaki and Draguhn, 2004) necessary for generation of preBötC population preI activity and I-bursts. To elaborate, the emergence of temporally aligned EPSPs and *V*_m_ correlation between I-M SST^+^ neurons (Figures 2A1-D, S2B1-B4) reflect the onset of output synchrony of their afferent rhythmogenic, including Type-1, neurons. The resultant synchronous firing of I-M SST^+^ neurons then leads to I-bursts (Figure 2E), with a subsequent decrease in input synchrony resulting in termination of the I-burst.

### GABA_A_ inhibition regulates preBötC synchronization and conductance state of I-M SST^+^ neurons

What network mechanism(s) leads to the emergence of preBötC synchrony that appears to underlie rhythmicity? Evidently, an overall (uncorrelated) increase in the neuronal firing is insufficient (Figures 2G-I, S2D1-E). Given that the excitation-inhibition balance is a critical determinant of network output (Bartos et al., 2002), including for breathing (Baertsch et al., 2018; Baertsch et al., 2019; Janczewski et al., 2013; Xiaolu Sun, 2019), we hypothesized that the net impact of elevated [K^+^]_ACSF_ is to shift the excitation-inhibition balance towards higher excitation, thus favoring preBötC synchronization. To test this hypothesis, we disinhibited the preBötC network by antagonizing gamma-aminobutyric acid type A (GABA_A_) or glycinergic receptors. Surprisingly, antagonism of GABA_A_, but not glycine, receptors in a nonrhythmic slice resulted in rhythmic I-bursts associated with synchronous EPSPs in I-M SST^+^ neurons (3A-D, S3A-F). Strikingly, for I-M SST^+^ neurons, at the onset of rhythmic bursting, despite the emergence of input synchrony, *V*_m_ global wavelet power decreased, as did EPSP amplitude, durations, and 20%–80% rise times (Figure 3E, F-H). In a subset of these experiments, we continuously monitored input resistance (*R*_in_) and membrane time constant (*τ*_m_) (N=6 neurons from 6 slices) during the transition from nonrhythmicity to the emergence of input synchrony and rhythmicity in response to GABA_A_R blockade, and found that values for both decreased (Figure 3I-J). This shift indicates onset of a higher conductance (HC) state.

**Figure 3.**
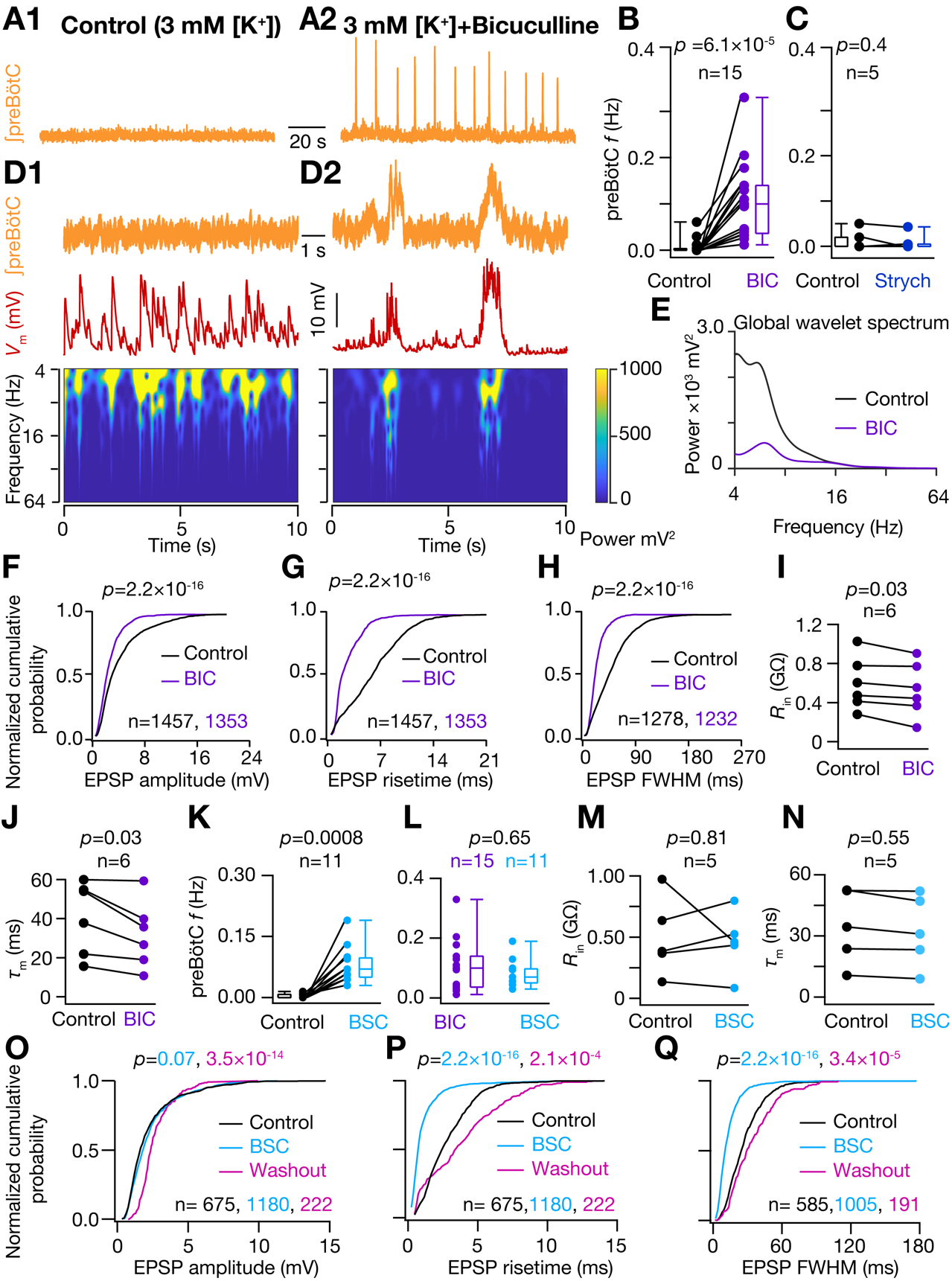
GABA_A_ inhibition regulates preBötC synchronization and conductance state of I-M SST^+^ neurons. (A) preBötC activity under nonrhythmic control (A1) and under 10 μM Bicuculline (BIC) rhythmic (A2) conditions. (B-C) preBötC burst frequency under control and BIC (B) and 2 μM strychnine (Strych) (C) conditions. (D) frequency-time plot for *V*_m_ of I-M SST^+^ neuron under control (D1) and BIC rhythmic (D2) conditions. (E) global wavelet spectrum of *V*_m_ in (D1-D2), note decrease in global wavelet power under rhythmic conditions with BIC. (F-H) normalized cumulative probability for spontaneous EPSP amplitude (F), 20%-80% rise time (G), and full width at half maximum duration, FWHM (H), under control (black) and rhythmic with BIC in ACSF (purple) conditions; N=8 neurons from 8 brain slices. (I) Input resistance, *R*_in_, and (J) membrane time constant, *τ*_m_, of I-M SST^+^ neurons recorded in control and BIC. (K) preBötC burst frequency recorded under control and with 10 μM Bicuculline, 2 μM Strychnine and 2 μM CGP55845 (cocktail abbreviated as BSC) in ACSF to block GABA_A_, glycinergic and GABA_B_ receptors, respectively. (L) comparison of preBötC frequency under control, BIC and BSC conditions. (M-N) *R*_in_ (M) and *τ*_m_ (N) of I-M SST^+^ neurons recorded in control and BSC conditions. (O-Q) same as (H-J) respectively, but comparing control (black), BSC (cyan) and after washout of BSC with control ACSF (pink); *p* values for comparison of color-coded experimental sets vs. control; N=5 neurons from 5 brain slices. For (B-C), (I-K) and (M-N), *p*-values are for Wilcoxon signed rank test; for (F-H), (L) and (O-Q), *p*-values for Wilcoxon rank sum test

The induction of a HC state in preBötC neurons by GABA_A_R blockade contrasts with the neocortex, where active inhibitory receptors are instead dominant determinants of a HC state (Destexhe et al., 2003). Therefore, we further explored roles of glycinergic and GABA_B_ receptors in effectuating HC state in preBötC I-M SST^+^ neurons. When three types of inhibitory amino acid receptors, i.e., GABA_A_, GABA_B_ and glycine, were all blocked in 3 mM [K^+^]_ACSF_, there was no significant difference in the preBötC frequency compared to GABA_A_R blockade only (Figure 3K-L), suggesting that under these conditions network synchronization and rhythmogenesis are exclusively modulated by GABA_A_-mediated inhibition. Furthermore, with blockade of all three inhibitory receptors, there was no significant change in either *R*_in_ or spontaneous EPSP amplitude (Figure 3M, O). The reduction in *τ*_m_ was also less compared to that during only GABA_A_R blockade (Figures 3N, S3I-J); there was still a significant decrease in EPSP rise time and duration (Figure 3P-Q), suggesting that integrative neuronal properties may be also regulated by excitatory synaptic and calcium-dependent intrinsic conductances under rhythmic conditions (Del Negro et al., 2010). Importantly, changes in EPSP rise time and duration were reversible (Figure 3P-Q), ruling out the possibility that these changes were primarily due to plasticity in neuronal response dynamics caused by increased cytosolic Ca^2+^ mobilization with increased network excitability (Ashhad et al., 2015; Ashhad and Narayanan, 2019; Narayanan et al., 2010). Taken together, these results uncover a significant and novel dichotomy in the role of GABAergic and glycinergic inhibition in breathing rhythmogenesis. While GABA_A_ regulates network synchrony, glycinergic and GABA_B_ inhibition each can affect mechanisms that shape pattern and assure propagation of I-bursts through modulation of synaptic integration and of the excitability of preBötC output neurons. Additionally, our results suggest that antagonizing GABA_A_ receptors disinhibits glycinergic inhibition onto I-M SST^+^ neurons, contributing to a HC state. These observations are direct evidence that network synchronization, irrespective of any significant increase in average population firing rate, is sufficient for preBötC rhythmicity (Figure 3D-E). Furthermore, I-M SST^+^ neurons can transition to a HC state and show rhythmicity in going from 3 mM to 9 mM [K^+^]_ACSF_, possibly as a consequence of increased activity of both excitatory and inhibitory neurons (Figure S4). Thus, the network synchronization and increased conductance state of these neurons are tightly coupled, representing an emergent feature of the preBötC microcircuit common across experimental conditions employed to induce rhythmicity.

### Propagation of preBötC bursts to XIIn is dependent upon strength of input synchrony onto I-M SST^+^ neurons

Given the central role of network synchronization in generation of preBötC I-bursts, we explored its role in their efferent propagation to various motor nuclei that control inspiratory muscles. We noticed that under rhythmic conditions ~16% I-M SST^+^ neurons (4/25 neurons recorded in 9 mM [K^+^]_ACSF_) received more bouts of synchronous inputs than the number of I-bursts that propagated through XIIn in the same time frame (Figures 1B2, 2E blue trace, 2F, 4A-C). Thus, when the input synchrony was weak or short-lived (reduced area of significant increase in the *V*_m_ power in frequency time plot), preBötC burstlet activity (Kam et al., 2013a) did not propagate to the XIIn. This observation is akin to a synfire chain, where propagation of synchronous activity along successive neuronal populations is governed by temporal compactness of population activity (here due to synchrony) and total number of spiking neurons in each population (Diesmann et al., 1999; Kumar et al., 2010). Under such an arrangement, the output synchrony of a given network serves as input synchrony for neurons in the downstream network, so that the stable propagation of population bursts along a linear network is a consequence of its input synchrony and total activity of its afferent network (Diesmann et al., 1999). In support of this theory, we found that when the input synchrony onto I-M SST^+^ neurons was weak, preBötC I-burst activity either failed to initiate or was truncated as a burstlet (Figure 4B-C). Additionally, bouts of synchronous EPSPs associated with XIIn I-bursts were sustained for significantly longer compared to those bouts that failed to produce XIIn I-bursts (Figure 4D-G). Furthermore, successful propagation of preBötC activity was codependent on both the duration of input synchrony and the resultant amplitude of *V*_m_ deflections (Figure 4H), which is a function of temporal compactness as well total activity of the presynaptic population.

**Figure 4.**
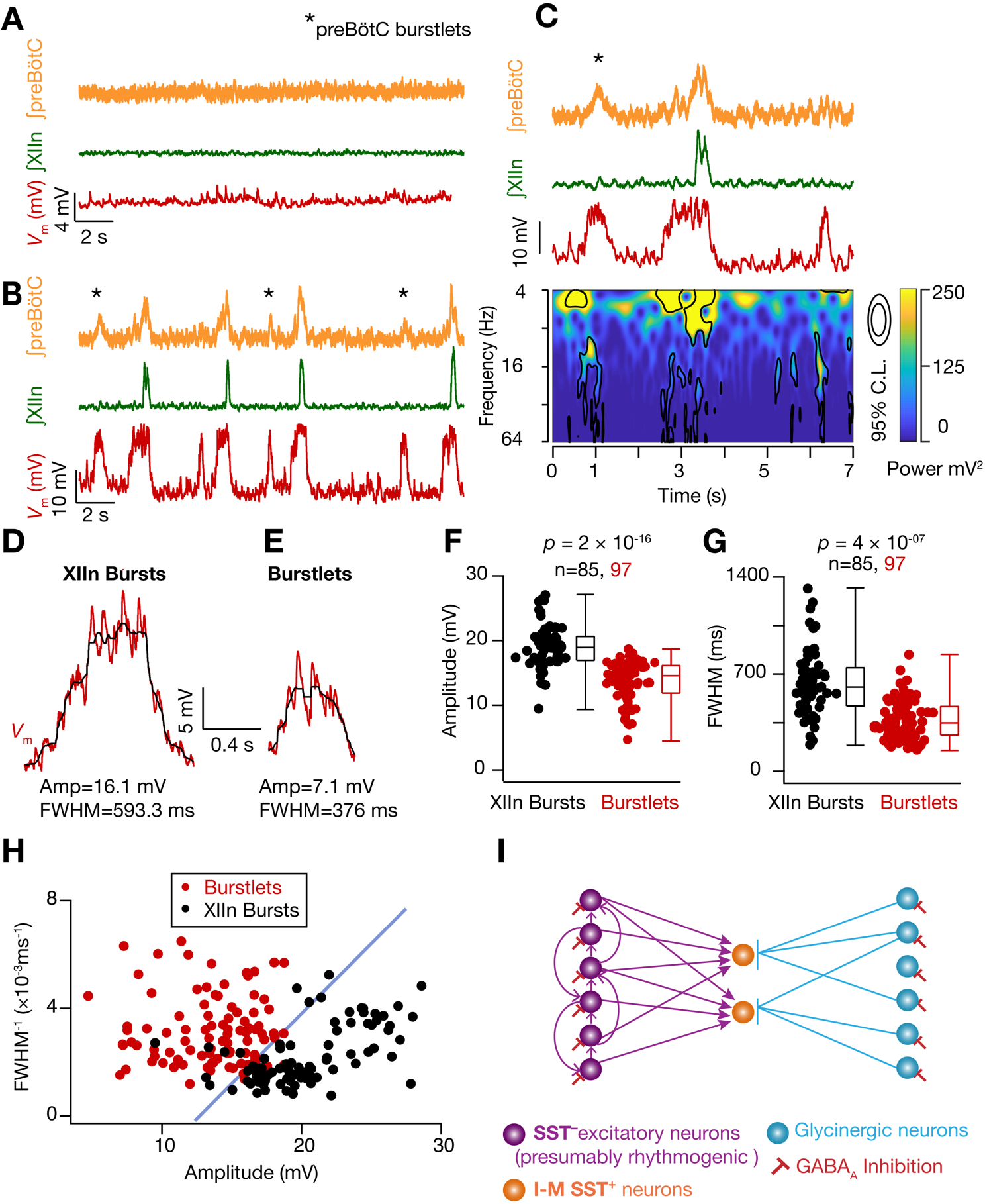
Propagation of preBötC bursts to XIIn is dependent upon strength of input synchrony onto I-M SST^+^ neurons. (A-B) preBötC and XIIn activity and *V*_m_ of an I-M SST^+^ neuron under control (A) and rhythmic (9 mM [K^+^]_ACSF_) (B) conditions. Note more bouts of input synchrony in *V*_m_ than preBötC and/or XIIn I-bursts in B, i.e., burstlets. (C) frequency-time plot of first 7 seconds from (B). (D-E) representative *V*_m_ from another neuron during an I–burst (D) and when preBötC input synchrony was not accompanied by an I–burst in XIIn (E). Raw (red) and median filtered (black) *V*_m_; later was used to compute amplitude and FWHM of *V*_m_ deflections during bouts of input synchrony. (F-G) summary plots for measurements in (D-E); n = number of events. (H) plot of FWHM^−1^ vs amplitude of *V*_m_ deflections from I-M SST^+^ neurons (N=4) during bouts of input synchrony. The data is color coded for synchronous inputs that resulted in XIIn I-bursts (black) and those that did not (red). An arbitrary blue line with slope of 1 ms x mV separates the data into the two groups. (I) Schematic representation of inputs onto I-M SST^+^ neurons as inferred from this study. These neurons receive synchronized inputs from SST^−^ glutamatergic neurons, some of which are presumptively rhythmogenic and connected through excitatory synapses among themselves (Rekling et al., 2000). I-M SST^+^ neurons also receive glycinergic inhibition that is also regulated by GABA_A_ inhibition.

## DISCUSSION

Delineating essential features of network dynamics underlying inspiratory rhythmogenesis has been a critical missing link in the endeavor to understand the neural basis of breathing behavior. Here, exploiting a unique experimental preparation where the rhythmogenic kernel is well-defined and propagation of rhythmic activity can be tracked to a motor nerve, and through direct comparison of network activity under nonrhythmic and rhythmic conditions, we found that network synchronization is essential for rhythmicity and its propagation to motor nerves (Figure 4I). While synchronous inputs reliably drive each inspiratory cycle, the underlying evolution of the EPSP correlation between I-M SST^+^ neurons varies on a cycle to cycle basis (Figure 2B1-B3), implying that the detailed network dynamics vary, i.e., in each cycle the network synchronizes through reliable formation of dynamical cell assemblies (Buzsaki and Draguhn, 2004; Carroll and Ramirez, 2013; Tsukada et al., 1996) that has a random or chaotic element. The inherent variability in the evolution of synchrony in this microcircuit could manifest as jitter in preI duration (which marks the onset of network synchronization) observed in slices (240 ± 10 ms; mean ± sem; range = 63 ms to 588 ms; data for preI events in Figure 2B1) (also compare Figures 1C3, 2A and 4C), as well as in the preceding IBIs (which reflects the time necessary for the network to (re)assemble). Additionally, the preI duration mirrors the latency to induce ectopic I-bursts following synchronous stimulation of very few (3-9) randomly chosen preBötC neurons *in vitro* (Kam et al., 2013b); we hypothesize that this latency to I-burst onset reflects the same process as underlying preI, where synchronization is seeded by random concurrence in activity of a few neurons in the preBötC rhythmogenic microcircuit. Similarly, under conditions of reduced network drive, cycle-to-cycle temporal variability in the evolution of network synchrony can contribute to increased jitter in the period of I-bursts, i.e., breathing frequency (Wang et al., 2014) (Figure S3A-B).

Exploring network mechanisms underlying synchrony, our results indicate that activity of GABAergic neurons regulate synchronization of rhythmogenic (SST^−^) preBötC neurons. Thus, under control conditions, active GABA_A_ inhibition can prevent synchronization of these neurons. Once GABA_A_Rs were blocked, these neurons could synchronize periodically to give rise to rhythmic inspiratory motor command. Counterintuitively, even when the inhibition was substantially reduced (by GABA_A_R blockade) the overall synaptic activity onto I-M SST^+^ neurons decreased during IBI (Figures 3D, S3 G-H). Consequently, global wavelet power of the *V*_m_ of I-M SST^+^ neurons also decreased (Figure 3E). Thus, the emergence of periodic input synchrony onto I-M SST^+^ neurons along with a decrease in the input frequency during IBI, with GABA_A_R blockade, suggests that the effect of GABA_A_R blockade onto these neurons is not merely gain modulation. Rather, GABA_A_ inhibition is a critical regulator an emergent network dynamics within preBötC. Antagonizing GABA_A_Rs also led to a HC state in I-M SST^+^ neurons that was abolished, at least in part, upon blockade of glycinergic and GABA_B_ receptors. This suggests that blocking GABA_A_R in preBötC can result in increased activation of glycinergic neurons, possibly as a consequence of disinhibition of these neurons. As the HC state resulted in a decrease in EPSP amplitude (Figure 3F), this could also contribute to the decrease in spectral power in the *V*_m_ of I-M SST^+^ neurons upon GABA_A_R blockade (Figure 3E). Taken together, these experiments reveal complex interactions among various neuronal subtypes underlying the switch from an asynchronous and nonrhythmic state to a synchronous and rhythmic state of preBötC (Figure 4I).

Network synchronization and HC state in the output I-M SST^+^ neurons concurrently interact to produce rhythmic bursts. This is a manifestation of efficient cellular and circuit level coordination (Barlow, 1961; Das and Narayanan, 2017; Deneve et al., 2017). The HC state changes neuronal integrative properties such that a shorter *τ*_m_ and an increased temporal resolution renders them more responsive to higher frequency inputs, thereby favoring coincidence detection and enhancing their selectivity to synchronous inputs (Contreras and Steriade, 1996; Destexhe et al., 2003; Fernandez et al., 2011; Ratte et al., 2013; Rudolph and Destexhe, 2003; Stevens and Zador, 1998). This selectivity, especially during preI and I-bursts, will facilitate propagation of synchronous population activity in the downstream efferent neuronal populations (Diesmann et al., 1999; Kumar et al., 2010; Ratte et al., 2013). Thus, synchrony propagation in the inspiratory motor network (Figure S5) is seen in medullary inspiratory premotoneurons (Mitchell and Herbert, 1974) projecting onto hypoglossal (Wang et al., 2002) and phrenic (Parkis et al., 2003) motoneurons, as well as in the high frequency (HFO; 50-100 Hz) and medium frequency (MFO; 20-50 Hz) oscillations present during inspiration in the: i) discharge patterns of inspiratory premotoneurons and motoneurons (Christakos et al., 1991; Ellenberger et al., 1990; Feldman et al., 1980; Huang et al., 1996; Tan et al., 2010; Yang and Feldman, 2018), and; ii) motor nerves projecting to inspiratory muscles (Christakos et al., 1991; Funk and Parkis, 2002; Schmid et al., 1990). Taken together, strong association between the network dynamics and neuronal properties that facilitate synchrony transfer (Ratte et al., 2013) is compelling evidence that the inspiratory motor command is a form of temporal neuronal code (Buzsaki and Draguhn, 2004; Srivastava et al., 2017), which, among other possibilities, increases the reliability of propagation of inspiratory drive from its source, i.e., preBötC, to the ultimate neural output, i.e., motoneurons.

While our study focuses on mechanisms of inspiratory rhythmogenesis, I-burst initiation and propagation, the mechanisms of I-burst termination and control of IBI remain open questions. Both burstlets and I-bursts self-terminate, even when all inhibition is blocked (Figure 3K,L). We speculate that this is due to a decrease in synaptic efficacy following network synchronization, e.g., due to increased neuronal conductance (Bernander et al., 1994). Moreover, while not essential to rhythmogenesis, inhibition originating in the preBötC does regulate the duration and shape of I-bursts, and modulates breathing frequency (Baertsch et al., 2018; Baertsch et al., 2019; Janczewski et al., 2013; Shao and Feldman, 1997; Sherman et al., 2015).

Finally, recent studies have uncovered widespread coupling of breathing rhythms with many other brain regions regulating or affecting behaviors, e.g., whisking (Moore et al., 2013), attention control (Yackle et al., 2017), and possibly higher cognitive function through bottom-up respiratory modulation of neuronal activity in hippocampus, and prefrontal cortex (Karalis N, 2018). In the light of divergent ascending projections of preBötC output neurons through mono- and oligo-synaptic connections (Yang and Feldman, 2018), synchronized oscillations in inspiratory motoneurons (Christakos et al., 1991; Funk and Parkis, 2002) and entrainment of limbic neurons by respiratory corollary discharge (Karalis N, 2018), we hypothesize that preBötC synchrony plays a critical role in the extraordinary reliability and robustness of networks underlying respiratory motor output as well as in the suprapontine propagation of breathing rhythms that may serve for binding of activity with and across higher brain regions.

The unique studies here, where we probed the mechanisms underlying a directly measurable output (motoneuronal activity) of a critical behavior, i.e., breathing rhythmogenesis, were possible because the preBötC is the only mammalian CPG with a clearly localized and well-defined kernel. We suggest that the mechanisms of synchrony revealed in this novel probing of neural circuits underlying breathing movements represent a fundamental principle of network signal processing in mammals underlying behavior.

## Acknowledgements

This work was supported by NIH-NHLBI grant 1R35HLI35779. We thank D.V.Buonomano, A.Kumar and R. Narayanan for their insights and discussion on this project and R. Abreu, D.N.Chiu, Hailan Hu, K.Kam, M.Shao and C.F.Yang for their comments on earlier versions of this manuscript.

## Author Contributions

S.A and J.L.F designed and interpreted experiments, S.A performed experiments and analyzed data. S.A. and J.L.F critically evaluated data, wrote paper and approved the final version of the manuscript.

## MATERIALS AND METHODS

### EXPERIMENTAL MODEL AND SUBJECT DETAILS

#### Mice

Animal use was in compliance with the guidelines approved by the UCLA Institutional Animal Care and Use Committee. Mice were housed in vivarium with 12-hour light/dark cycle and food and water was supplied ad *libitum*. Adult SST-Cre (IMSR Cat# JAX:013044, RRID: IMSR_JAX:013044) mice were crossed to Ai14 Cre-reporter mice (IMSR Cat# JAX:007914, RRID: IMSR_JAX:007914) to generate the SST reporter line used for *in vitro* slice experiments. All experiments were performed using the reporter line neonates between postnatal 0 to postnatal 7 days old.

### METHOD DETAILS

#### In Vitro slice preparation and electrophysiological recordings

The neuraxis containing the brainstem and the spinal cord, from neonatal mice of either sex, was isolated and one transverse brainstem slice (550 μm-600 μm thick) was prepared from each animal that contained the preBötC along with respiratory premotor and hypoglossal (XII) respiratory motor neurons and XII nerve (XIIn) rootlets. The slice was cut (using a Leica VT1200 slicer) such that the preBötC was on the surface of the slice when placed rostral side facing upwards and contained anatomical landmarks as described in(Ruangkittisakul et al., 2014). The slices were cut in chilled ACSF containing (in mM): 124 NaCl, 3KCl, 1 MgSO_4_, 25 NaHCO_3_, 0.5 NaH_2_PO_4_ and 30 D-glucose, bubbled with 95%CO_2_ and 5% O_2_, at 7.4 pH

For electrophysiological recordings, slices were perfused with ACSF (32-34°C) at 4 ml/min in the recording chamber where they were allowed to recover for at least 30 minutes in normal ACSF before the start of recordings. To record preBötC and XIIn population activity, ACSF filled suction electrodes were used (with ~150 μm tip diameter) pulled from borosilicate glass (1.2 mm outer diameter with 0.52 mm wall thickness) using P-97 puller from Sutter Instruments. Data was acquired through a differential AC amplifier (A-M Systems, Model 1700), filtered at 1-5 kHz and digitalized using a MultiClamp 700B (Molecular Devices). The sampling frequency was either 20 or 40 kHz. The population recording time series was full wave rectified and integrated using a custom built analog Paynter filter with a time constant of 15 ms. The integrated data lacked scale for comparisons across experiments and, thus, was represented as normalized arbitrary units(Kam et al., 2013a).

For intracellular whole-cell current-clamp clamp recordings, fluorescently labeled (tdTomato) somatostatin-positive (SST^+^) neurons were patched under visually guided video microscopy using an upright microscope (either Zeiss Examiner A1or Olympus BX51W1) fitted with DIC optics and CCD camera (Andor Technology iXon or Hamamatsu C11440). The preBötC boundary was ventral to nucleus ambiguous and lateral to the principle loop of inferior olive. Neurons were patched with glass electrodes of ~1μm tip diameter with electrode resistance of 5-8 MΩ. Electrodes were pulled from borosilicate glass capillary (1.5 mm outer diameter, 0.64 mm wall thickness) using P-97 microelectrode puller (Sutter Instruments) and filled with intracellular solution containing (in mM): 120 K-gluconate, 20 KCl,10 HEPES, 4 NaCl, 4 Mg-ATP,0.3 Na-GTP, and 7 K_2_-phosphocreatine (pH= 7.3 adjusted with KOH, osmolarity ~310 mOsm). Whole-cell current-clamp recordings were performed on SST^+^ neuronal somata using Dagan IX2-700 amplifier and data was digitalized using a MultiClamp 700B at either 20 or 40 kHz. Measured liquid junction potential (LJP) between the pipette solution and ACSF was 16-20 mV and it was not compensated during the experiment. Unless otherwise noted, all neurons were hyperpolarized to ~−75 mV by negative current injection to record synaptic inputs and minimize their AP firing. Resting membrane potential at break-in (without LJP compensation) was between −48 mV to −60 mV. Series resistance (*R*_s_) and series capacitance (*C*_s_) was monitored by examining the voltage response of the patched neurons to a hyperpolarizing current injection of 10 pA and was compensated online through bridge balance, though high synaptic noise and hence fluctuating membrane potential meant that the compensation was only approximate. *R*_s_ range was between 13 MΩ-40 MΩ.

#### Pharmacology

To block *γ*-aminobutyric acid (GABA) type A receptors (GABA_A_R), glycine receptors (GlyR) and GABA_B_ receptors 10 μM (−)Bicuculline methiodide (Tocris Bioscience), 2 μM Strychnine hydrochloride (Tocris Bioscience) and 2 μM CGP55845 (Tocris Bioscience) were used respectively.

To induce inspiratory rhythm in preBötC slices either [K^+^]_ACSF_ was increased to 9 mM or inhibition was pharmacologically blocked in normal, i.e., 3 mM [K^+^]_ACSF_.

#### Simulations

A biophysically realistic conductance based single-compartmental model of preBötC neuron was constructed as a cylinder 20 μm in diameter and 20 μm long with the following passive parameters: membrane capacitance, *C*_m_ = 1 μF/cm^2^; specific membrane resistance (*R*_m_) = 10 MΩ.cm^2^ and axial resistivity (*R*_a_) = 200 Ω.cm. These parameters were chosen such that the input resistance, *R*_in_, of the model neuron was 350 MΩ, which falls in the range of experimentally observed values of *R*_in_ for I-M SST^+^ neurons (Figures 3I and 3M). Two voltage-gated ion channels, i.e., a fast sodium (NaF) and a delayed rectifier potassium (KDR), were incorporated into the model. Each ion channel was modeled using a Hodgkin-Huxley formulation adopted from(Migliore et al., 1999). NaF channel density was 45 mS/cm^2^ and KDR channel density was 15 mS/cm^2^. Current through α-amino-3-hydroxy-5-methyl-4-isoxazolepropionic acid (AMPA) receptors was modeled as a combination of sodium and potassium currents:

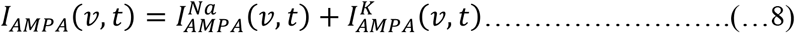

where,

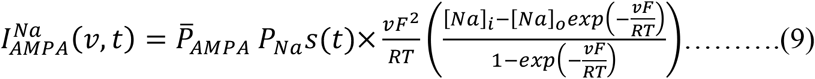

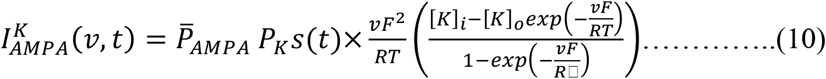

where 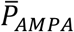 is the maximum permeability of AMPAR that was set to 1×10^−7^ cm/s so that the resultant EPSP amplitude was 2.5 mV, which approximates the mean amplitude of spontaneous EPSPs of I-M SST^+^ neurons under normal ACSF (Figures 3F and 3O). *P*_Na_ was set equal to *P*_K_=1 (Dingledine et al., 1999). *s*(*t*) determined the kinetics of current through AMPAR and was defined as:

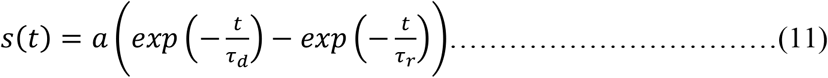

where *a* is a normalization constant to ensure that 0≤*s*(*t*)≤1. *τ*_r_=2 ms and *τ*_d_=10 ms(Narayanan and Johnston, 2010).

All simulations were performed in the *NEURON* simulation environment(Hines and Carnevale, 1997) with the integration time step of 25 μs. Resting membrane potential was set at −75 mV, simulation temperature was 34°C and ion channel kinetics were adjusted per their experimentally determined *Q*_10_ coefficients.

#### Data analysis

To estimate input resistance, *R*_in_, a hyperpolarizing pulse of 10 pA current pulse of 1s duration was applied for 10-30 trials (Figure S3G-H). The average membrane potential deflection was then divided by the injected current to get the estimate of *R*_in_. To compute membrane time constant, *τ*_m_, the repolarizing phase of the averaged *V*_m_ trace was fitted with an exponential function of form:

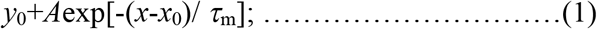

where y_0_, A and *τ*_m_ are fit coefficients and x_0_ is the steady-state value of *V*_m_ at the end of repolarization. For these estimates, only trials in the interburst intervals were considered.

To compute population firing rate from population recordings (Figure S1 A3) raw signal was first band-pass filtered between 300 Hz to 3 kHz to get multiunit activity (MUA). From this MUA, extracellular action potentials, exhibiting positive and negative polarity, were detected as peaks that were greater than a predetermined threshold in either direction and were separated from each other by at least 1.5 ms (to avoid multiple detection of same spike). The threshold for spike detection was determined by the noise level (representing neuronal + signal noise) present under the non-rhythmic condition (in normal ACSF) and in this case (Figure S1A3) was set to 10 μV. Spike times of detected spikes was converted into instantaneous firing frequency which was then box smoothened twice, with a window of 2 ms, to give rise to the final population firing rate trace.

#### Wavelet analysis

Time frequency decomposition of neuronal *V*_m_ by continuous wavelet transform (CWT) was implemented through guidelines presented in(Torrence and Compo, 1998). Briefly, *V*_m_ was subtracted from its mean and normalized by its standard deviation. Thereafter, CWT was performed on the resultant time series, *x*_n_, defined as the convolution of *x*_n_ with scaled and translated version of the wavelet function Ψ_0_(η):

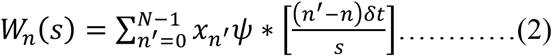

where the (*) indicates complex conjugate. The minimum scale, *s*_0_, was fixed at 0.0153 s (corresponding to 64 Hz). Subsequent scales were determined as:

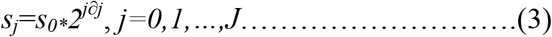

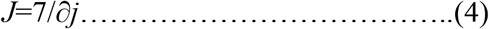

Where, *J* determined the largest scale and *∂j*, i.e., increment of *J*, was fixed at 0.025. Thus, the total number of scales was *J*+1= 281. Ψ is the normalized version of Ψ_0_(η) which is the complex Morlet wavelet defined as:

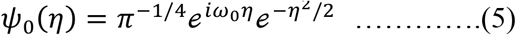

Where ω_0_ is the nondimensional frequency chosen to be 6 to satisfy the admissibility criterion for a Morlet wavelet(Torrence and Compo, 1998). η is the nondimensional time parameter. Ψ_0_(η) was normalized to give Ψ so that at each scale, s, the wavelet function has unit energy irrespective of the scale size:

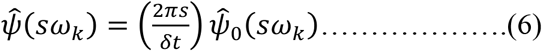

This made it possible to directly compare the wavelet transform at each scale and to the wavelet transforms of other time series(Torrence and Compo, 1998), i.e., wavelet transforms of *V*_m_ of a given neuron under inspiratory rhythmic and non-rhythmic conditions. The global wavelet spectrum was computed as the time averaged wavelet spectrum over entire time multiplied by the variance of the original signal. For significance testing the global wavelet spectrum of *V*_m_ under non-rhythmic condition (normal ACSF) was chosen as background spectrum for each neuron. Peaks in the wavelet spectrum of the *V*_m_ of the same neuron under rhythmic condition was then considered to be above the background spectrum with 5% significance level (or 95% confidence level) if it was above the product of background spectrum with 95^th^ percentile value for chi-square distribution having two degrees of freedom(Torrence and Compo, 1998). Wavelet software was provided by C. Torrence and G. Compo: http://paos.colorado.edu/research/wavelets/.

Unless otherwise stated, only epochs of *V*_m_ without any APs were used in the analysis. When *V*_m_ containing APs was used, they were removed through median filtering in a two-step process. First, in a large window of 10-13 ms all data points greater than 3 mV from the median were replaced by the median. This selectively removed AP peaks with sharp transients at the base of the truncated APs. The filtered signal was again median filtered with a 2 ms window to remove these transients. This process ensured that only APs were removed from the *V*_m_ trace without distorting synaptic potentials. To get the envelop of *V*_m_ deflections during the epochs of synchronous inputs (Figure 4 D-H) the traces were median filtered with a 25 ms window.

For paired whole-cell patch-clamp recordings, fluorescently labelled SST^+^ neuronal pairs were patched at intersomatic distance of 10-100 μm. The normalized cross-correlation between two time series, *x(t)* and *y(t)* (*V*_m_ of paired neurons for our analysis), was calculated as follows(Graupner and Reyes, 2013):

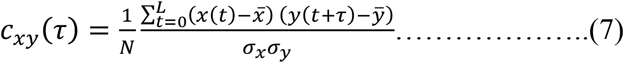

where, *L* is the total length of the signal, 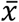 and 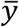 and *σ*_*x*_ and *σ*_*y*_ are means and standard deviations of *x(t)* and *y(t)* respectively. The value of *c*_*xy*_(*τ*) varies from −1 to 1 such that two identical signals will have a value of 1 at zero time lag (*τ* = 0). PreI duration was determined as the time between the onset of preBötC activity and the onset of an I-burst in the XIIn. The onset of preBötC activity was determined by the start of positive inflection in the differentiated ∫preBötC trace. For one I-M SST^+^ pair XIIn was not active, hence the preI period was determined by the initial slope of ∫preBötC trace (Figure S2 B4) (Kam et al., 2013a). Epochs of inter burst intervals (IBI) extracted from *V*_m_ trace were ~1 second after the termination of the I-burst given that they did not overlap with preBötC burstlets. For direct comparison, the duration of IBI epochs was fixed at the duration of the preI period preceding it.

To probe synaptic connections between the recorded neurons, a depolarizing current pulse was injected sequentially in each neuron to produce APs. The spike triggered average of *V*_m_ in non-firing neurons was computed to ascertain synaptic connectivity.

For quantification of spontaneous excitatory postsynaptic potentials (EPSPs), 100-300 s epochs of *V*_m_ from I-M SST^+^ neurons were used. Custom semi-automated peak detection software was used to select for spontaneous EPSPs. Due to very high synaptic noise, the start and end of each EPSP was manually specified to avoid temporally clustered EPSPs being falsely detected as a unitary EPSP. Only those EPSPs which did not have any visually detectable inflections in their rising phase were considered for analysis. When there were overriding EPSPs in the decay phase of a selected EPSP, its full width at half maximal (FWHM) duration was not computed. This resulted in a reduced number of events that were analyzed for FWHM as compared to EPSPs amplitude and rise time (*cf.* Figures 3 F-H and 3 O-Q). EPSP amplitude was defined as the difference of peak *V*_m_ from the baseline value for each event. 20%-80% rise time was defined as the duration in which the EPSP grew from 20% to 80% of peak amplitude. For non-I-M SST^+^ neurons, the recording duration was shorter and hence 50-100 s epochs were used for EPSP analysis.

## QUANTIFICATION AND STATISTICAL ANALYSIS

Data was analyzed using custom software in Igor Pro (version 7.4; WaveMetrics Inc.). Statistical analyses were performed using R software (www.r-project.org). Sample size was not predetermined. Normality was neither tested nor assumed and hence, for statistical testing, non-parametric tests were used. For box-and-whisker plots, center line represents the median; box limits, upper and lower quartiles and whiskers represent 90 and 10 percentile range.

## DATA AND CODE AVAILABILITY

The datasets generated during this study are available from the corresponding author upon reasonable request.

## Supplementary Figures

**Figure S1.**
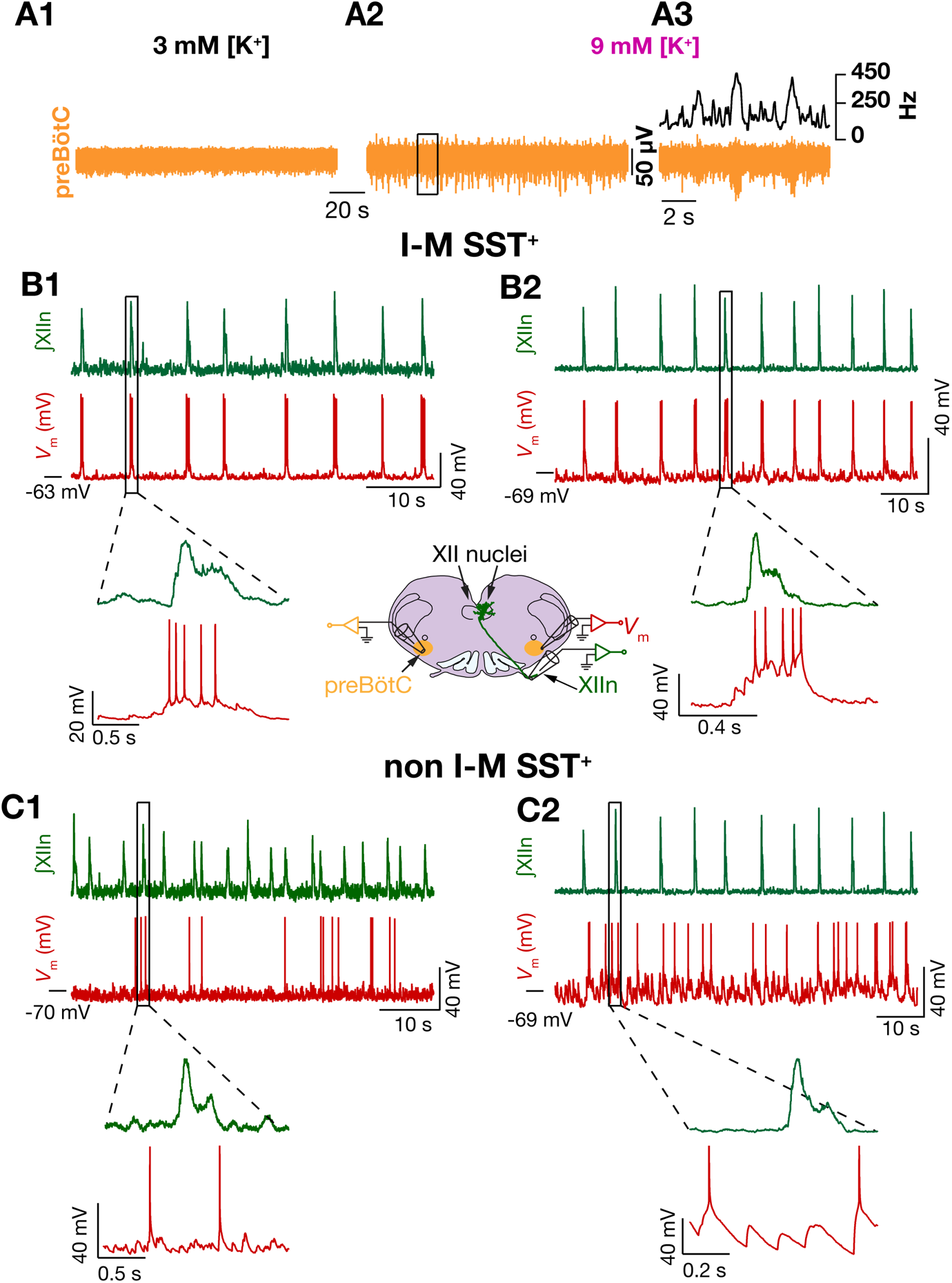
I-M SST^+^ neurons fire in phase with XIIn I-burst. (A) preBötC population activity in 3 mM [K^+^]_ACSF_, i.e., control (A1), and in 9 mM [K^+^]_ACSF_ (A2); (A3) expanded boxed region from (A2), top black trace is the instantaneous preBötC population firing frequency. These traces correspond to the integrated traces presented in Figure 1 A1-A3 respectively. (B1-C2) Firing profiles of I-M SST^+^ (B1-B2) and non I-M SST^+^ (C1-C2) neurons in 9 mM [K^+^]_ACSF_. Cartoon (center) of configuration for simultaneous recording of SST^+^ neurons in preBötC, XIIn and contralateral preBötC population activity. Traces in B1 are for the same neuron as in Fig. 1(B1-B3). Neurons in (B2) and (C2) were simultaneously

**Figure S2.**
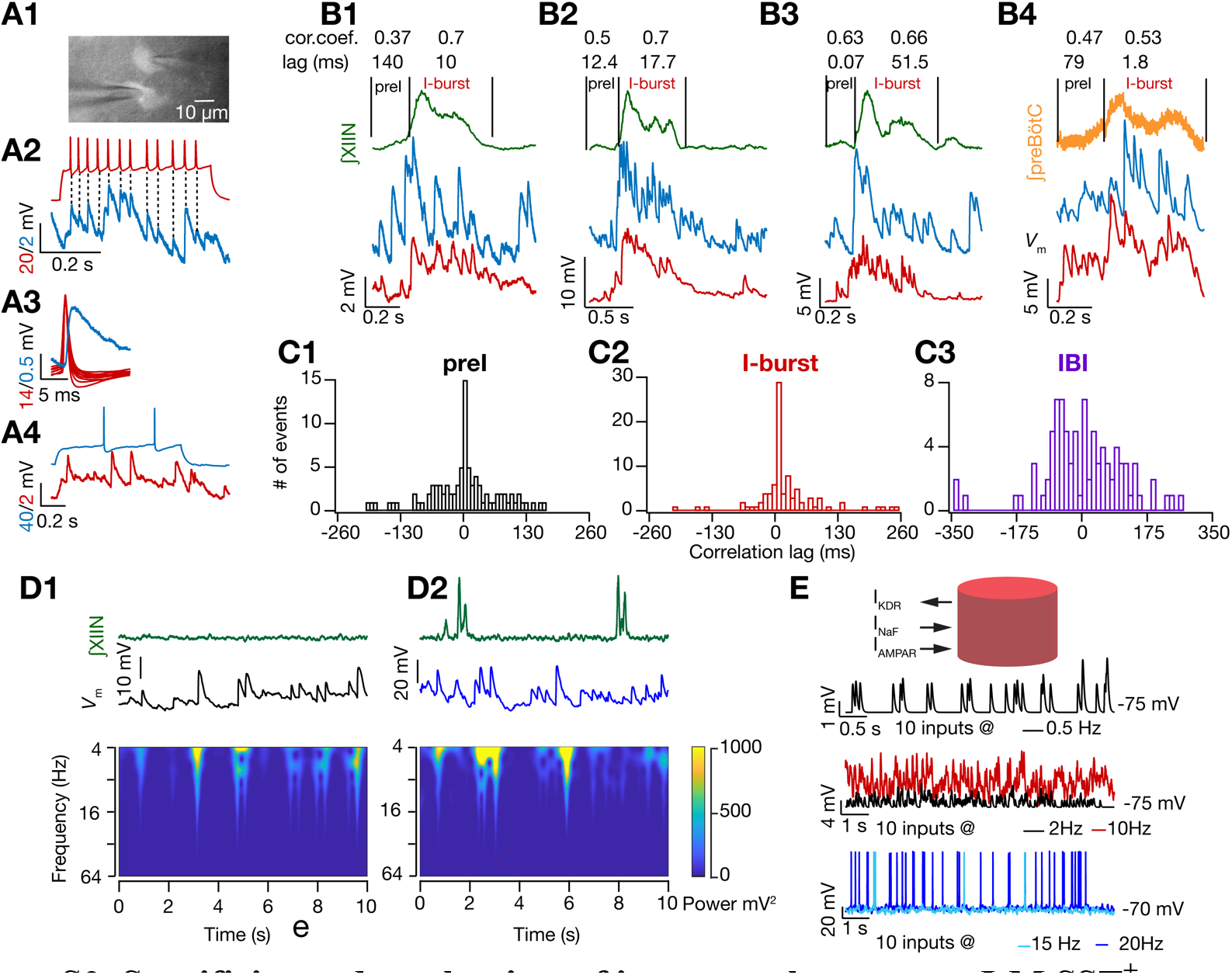
Specificity and mechanism of input synchrony onto I-M SST^+^ neurons. (A1) Fluorescent micrograph of simultaneously patched SST^+^ neurons. (A2-A4) *V*_m_ of a synaptically connected pair, represented by red and blue traces, when either of them depolarized to fire action potentials. (A3) spike triggered average *V*_m_ from (A2). The pair was unidirectionally connected with excitatory synapses, spike-triggered averaged *V*_m_ for the connected pair is also shown (middle); EPSP amplitude = 1.7 mV, latency to peak = 3.4 ms. This was the *only* synaptically connected pair among 50 pairs of SST^+^ neurons tested. (B1-B4) *V*_m_ of four simultaneously recorded I-M SST^+^ pairs (red, blue) with correlated EPSPs during preI and I-bursts. Traces in (A) and (B1) are from the same pair. Pairs (B2), (B3), (B4) were not synaptically connected. (C1-C3) Histogram of cross-correlation lags for *V*_m_ of I-M SST^+^ pairs for the data presented in Fig. 2 (B1-B3) where each point represents an “epoch”. (D1-D2) frequency-time plot of non I-M SST^+^ neuron same as in Figure. 2 G1-G2 but when the neuron was not spiking in 9 mM [K^+^]_ACSF_. (E) Representation of a single compartmental biophysical neuronal model with three currents, bottom: *V*_m_ under conditions where ten synaptic inputs were randomly activated at various frequencies as indicated.

**Figure S3.**
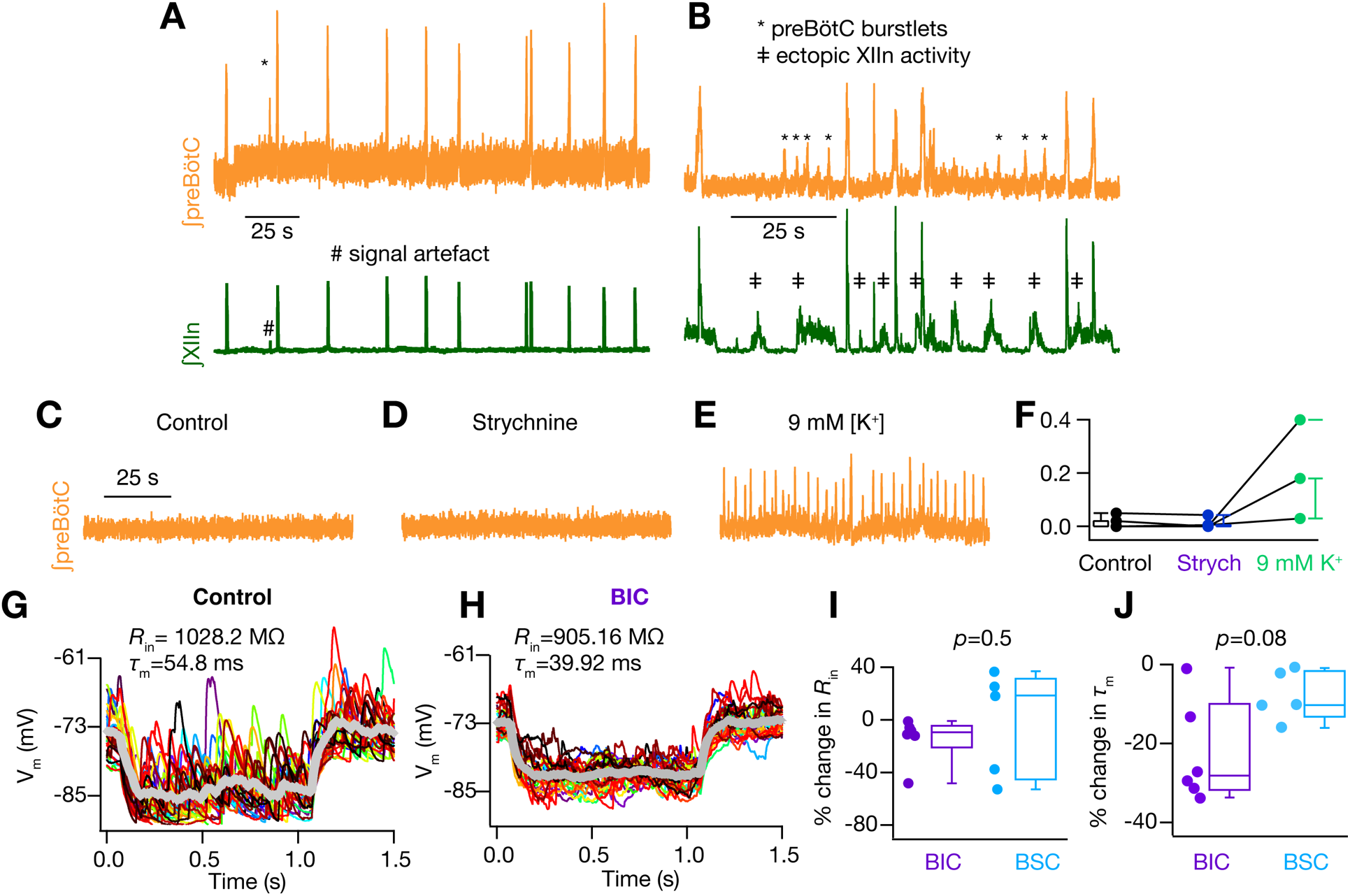
GABA_A_ inhibition gates preBötC rhythmogenesis and conductance state of I-M SST^+^ neurons. (A-B) simultaneously recorded preBötC (orange) and XIIn (green) activity in two brainstem slices showing small amplitude preBötC burstlets (marked by *) which did not result in XIIn I-bursts under 10 μM Bicuculline. In few slices, recorded under this condition, ectopic XIIn bursts ((B); marked ≠) were observed that were not concurrent with preBotC activity and, thus, were not considered for analysis. (C-E) preBötC activity of a representative slice recorded under control (C), 2 μM Strychnine (D) and in 9 mM [K^+^]_ACSF_ (E) after strychnine washout. (F) same as Fig. 3(C) but with additional experiments depicting activity of 3/5 slices that were also recorded in 9 mM [K^+^]_ACSF_ (green) after strychnine washout, e.g., as shown in (C-E). (G-H) *V*_m_ of an I-M SST^+^ neurons in response to a hyperpolarizing 10 pA current injection under control (G) and under 10 μM Bicuculline (BIC) (H); individual traces span 30 trials (depicted by different colors) and thick grey traces are averages under each condition. (I-J) % change in *R*_in_ (I) and *τ*_m_ (J) of I-M SST^+^ neurons recorded under BIC and BIC+Strychnine+CGP55845 (BSC) conditions; *p* values for Wilcoxon rank sum test.

**Figure S4.**
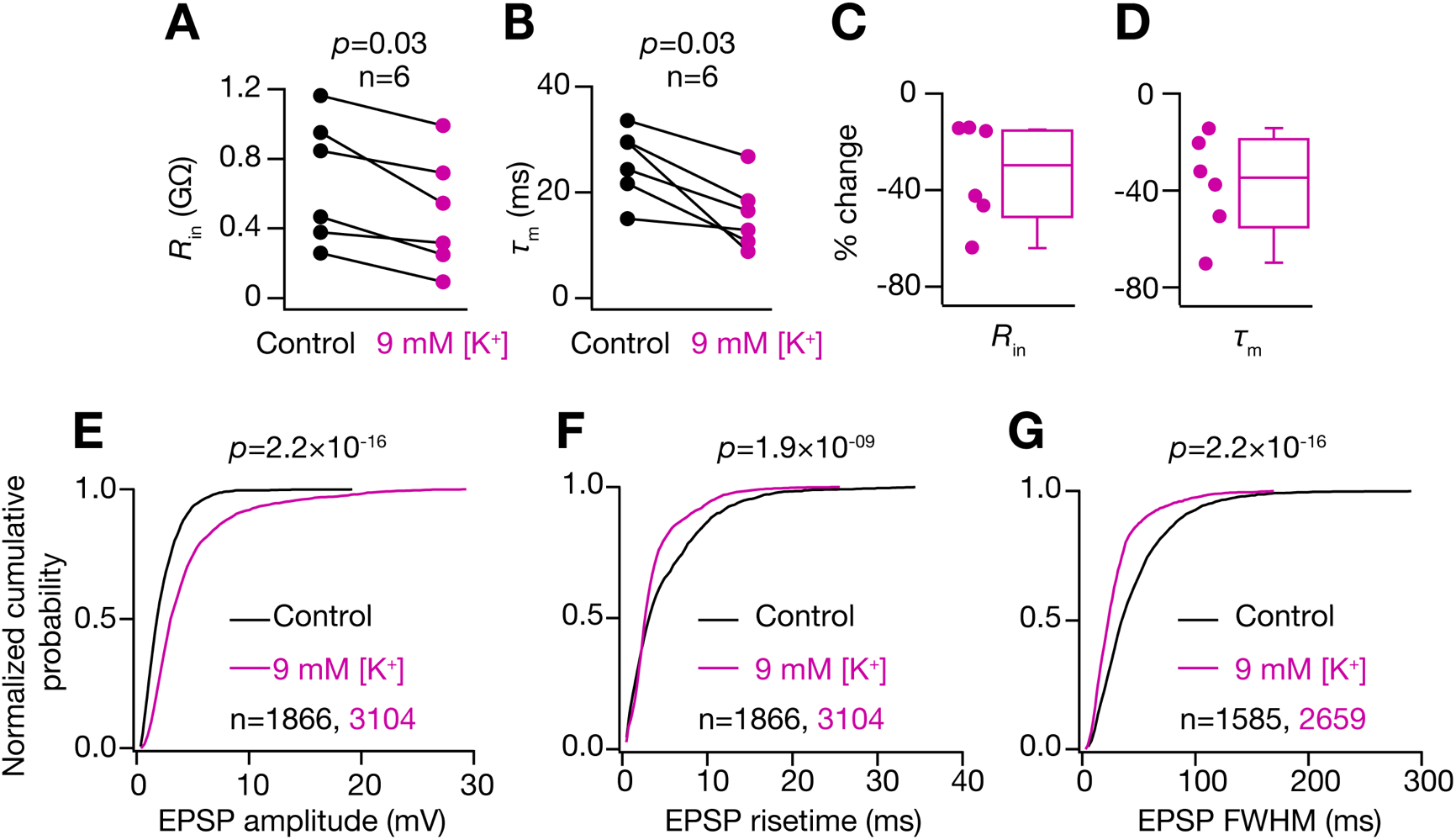
I-M SST^+^ neurons shift to higher conductance state in 9 mM [K^+^]_ACSF_. (A) *R*_in_, and (B) *τ*_m_, of I-M SST^+^ neurons recorded in control, i.e., 3 mM [K]_ACSF_, and rhythmic 9 mM [K^+^]_ACSF_ conditions. (C-D) percentage change in *R*_in_ (C) and *τ*_m_ (D) of these neurons under 9 mM [K^+^]_ACSF_. (E-G) normalized cumulative probability of spontaneous EPSP amplitude (E), 20%-80% rise time (F), and FWHM (G) of I-M SST^+^ neurons in control (black) and rhythmic 9 mM [K^+^]_ACSF_ (pink) conditions. Note the increase in EPSP amplitude in (E) even with a decrease in *R*_in_ of these neurons with change of ACSF from control to 9 mM [K^+^], which could be attributed to an increase in the presynaptic release probability with steady-state depolarization of axon terminals under 9 mM [K^+^]_ACSF_. (A-B) *p* value for Wilcoxon signed rank test; (E-G) *p* value for Wilcoxon rank sum test. For (E-G) N= number of neurons=14

**Figure S5.**
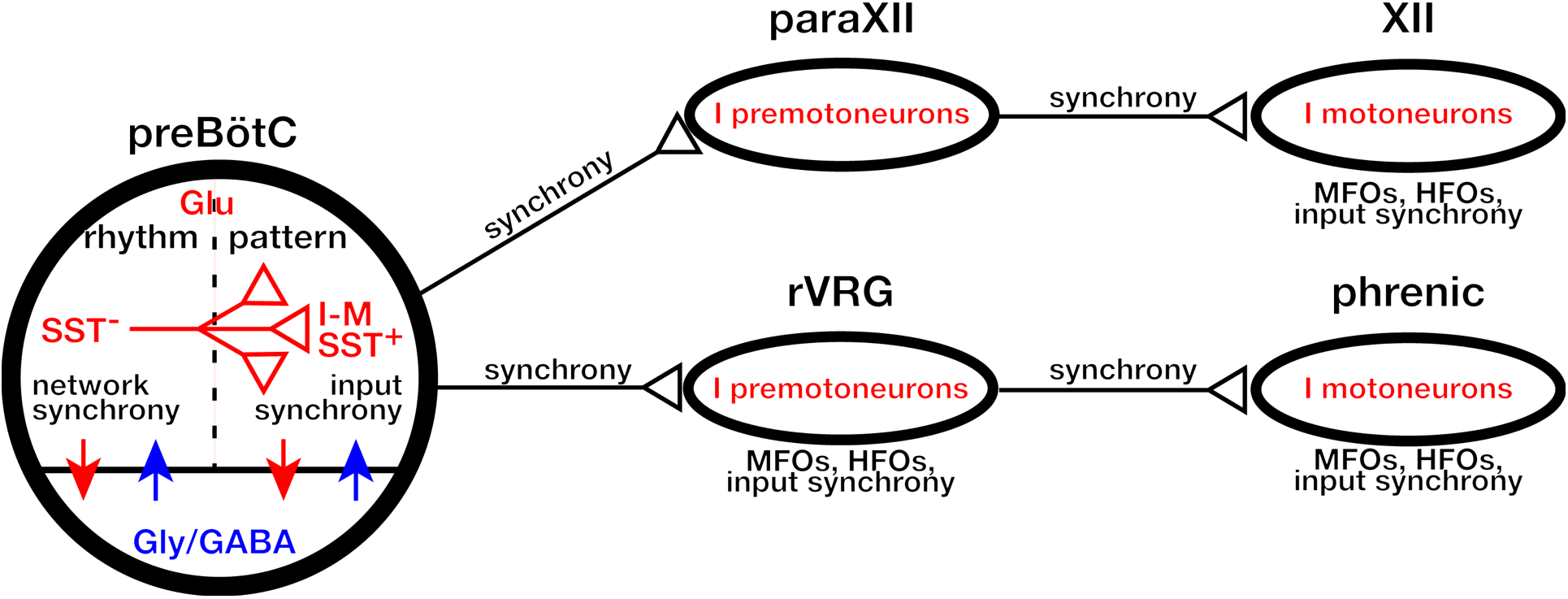
Synchronous propagation of preBötC activity to motor nuclei. Diagrammatic representation of population activity that emerges as synchrony in preBötC and propagates through inspiratory networks. Synchrony arises from correlated activity of preBötC rhythmogenic neurons and propagates via preBötC output neurons and premotoneurons to inspiratory motoneurons. MFOs = medium frequency (15-50 Hz) oscillations; HFO = High frequency (50-120 Hz) oscillations. Arrows indicate interactions between excitatory (red) and inhibitory (blue) neurons. The schematic is based on data presented in this paper as well as data represented in (Christakos et al., 1991; Ellenberger et al., 1990; Feldman et al., 1980; Funk and Parkis, 2002; Huang et al., 1996; Liu et al., 1990; Mitchell and Herbert, 1974; Parkis et al., 2003; Schmid et al., 1990; Tan et al., 2010; Wang et al., 2002; Yang and Feldman, 2018).

